# SpatialProp: tissue perturbation modeling with spatially resolved single-cell transcriptomics

**DOI:** 10.64898/2025.11.30.691355

**Authors:** Eric D. Sun, Alejandro Buendia, Anne Brunet, James Zou

## Abstract

Perturbational studies are the gold standard for identifying causal relationships between components of biological systems. Recent technological advances, including Perturb-seq and related assays, have enabled high-throughput screening of genetic perturbation effects on single cells. Several machine learning tools have also been developed to infer the effect of single-cell perturbations. However, both approaches are generally limited to dissociated cells, and the effect of genetic perturbations on neighboring cells within intact tissue has not yet been explored. Here we introduce a computational framework using graph neural networks for predicting the effect of multi-gene, multi-cell type perturbations on cells in whole tissue sections. We leverage the natural heterogeneity in tissue microenvironments across spatially resolved single-cell transcriptomics datasets to train SpatialProp (Spatial Propagation of Single-cell Perturbations). We show that SpatialProp can predict gene expression from the tissue microenvironment and map fine-grained steering of tissue microenvironments to new target states. To assess for causal enrichment in spatial perturbation predictions, we propose CausalInteractionBench, a bidirectional benchmarking approach using curated cell-cell interactions. Under this benchmark, we evaluate the causal utility of SpatialProp in predicting the spatial effects of different perturbations. SpatialProp provides a framework towards rapid hypothesis generation and *in silico* perturbation experiments, particularly in the study of spatially patterned tissue biology.

## Introduction

Perturbational approaches are key to building a causal understanding of cell and tissue biology. They exist in contrast to observational studies that report correlations or associations between different biological components, for example the expression of genes or abundance of proteins. A causal understanding of cellular and tissue systems is necessary to predict the dynamics of biological systems in response to perturbations or homeostatic imbalances, to disentangle compensatory and bystander effects, and to design targeted therapeutics for health and disease. Recent technological developments in functional genomics such as Perturb-seq and related approaches, which couple CRISPR genetic perturbations with transcriptomic readouts [1, 2, 3, 4, 5, 6], have enabled high-throughput studies of the effect of genetic perturbations on gene expression within single cells. However, these technologies harbor key limitations including cost and scalability, and they have largely been limited to dissociated cells rather than intact tissues. Several computational and machine learning models have been developed to bridge limitations of cost and scale with some success in modeling cell-intrinsic perturbations [ 7, 8,9, 10,11, 12, 13, 14, 15, 16, 17].

Importantly, there exists a technical gap between modeling the effect of single-cell perturbations in isolation and predicting how those perturbations influence neighboring cells in intact tissues. Single cells interact within a tissue to give rise to emergent functions that are relevant to complex processes like aging and disease. As such, there is a need to bridge our understanding of the first-order effects of genetic perturbations on an individual cell to the second-order effects of those perturbations on neighboring cells in the surrounding tissue microenvironment. Some recent efforts have extended single-cell perturbation studies to the spatial tissue context, including for transcriptomic measurements [18, 19, 20]. However, these technologies are still nascent and limited in their ability to resolve changes in spatial transcriptomic signatures at scale. Computational prediction of the effects of single-cell perturbations on a tissue microenvironment would enable flexible, high-throughput computational screens to identify promising candidate genetic interventions for many different applications that can then be validated more carefully through biological experiments. Early approaches in this direction have been applied to T cell perturbations in a tumor infiltration context using spatial proteomics [21] and for cell type transformations using spatial transcriptomics foundation models [22] and in tissue microenvironments [23]. A general-purpose framework, such as one that leverages gene expression across any number of cell types in a given tissue, will have broad utility across many different areas of tissue biology including in the contexts of rejuvenation, regeneration, and disease.

Here we introduce SpatialProp (Spatial Propagation of Single-cell Perturbations), which is a computational framework leveraging graph deep learning to predict the spatial effects of single-cell genetic perturbations using spatially resolved single-cell transcriptomics data. SpatialProp incorporates a simple graph neural network trained to predict center cell gene expression from gene expression data of neighboring cells (tissue microenvironment), leveraging natural heterogeneity in tissue microenvironments across spatially resolved single-cell transcriptomics data of intact tissues. SpatialProp uses this graph neural network within a calibration framework to propagate the effect of genetic perturbations in the tissue microenvironment to each cell, generating a perturbed tissue dataset in a cell-by-cell manner. We further introduce several evaluation frameworks for spatial perturbation models, which are models that predict the spatial effects of single-cell perturbations across a tissue. Our evaluation frameworks are designed to assess whether a given spatial perturbation model can reliably predict the gene expression of cells from their local microenvironment in the context of intact tissues, predict perturbed gene expression with fine-grained resolution sufficient to resolve intra-cell type heterogeneity (i.e. heterogeneity between cells of the same cell type), and produce predictions of spatial perturbation effects that are causally enriched for biological relevance (CausalInteractionBench). Together, SpatialProp and its associated benchmarks lay the groundwork towards computational simulation of spatial biology in response to complex single-cell perturbations.

## Results

### Spatial propagation of single-cell perturbations with SpatialProp

We introduce SpatialProp (Spatial Propagation of Single-cell Perturbations), which is a computational framework for predicting the spatial effects of single-cell genetic perturbations in intact tissues using spatially resolved single-cell transcriptomics data (Fig. 1A). SpatialProp consists of a simple graph neural network module, which is trained to predict gene expression of the center cell from the gene expression of neighboring cells in the surrounding tissue microenvironment, and additional modules for post-processing the predictions to generate perturbed gene expression. SpatialProp takes as input: a spatially resolved single-cell transcriptomics dataset of intact tissue and a user-defined set of single-cell perturbations represented by their perturbed gene expression profiles. Then, using the core graph neural network module, Spatial-Prop predicts perturbed gene expression in a cell-by-cell manner and calibrates these predictions for model error to update gene expression profiles for every cell in the tissue. Finally, SpatialProp outputs a prediction of the perturbed gene expression profiles for all cells in the spatially resolved single-cell transcriptomics data, including for cells that did not receive a direct user-specified perturbation. To assess the performance of SpatialProp and future spatial perturbation models, we further introduce several evaluation frameworks including CausalInteractionBench (Fig. 1B).

**Figure 1:**
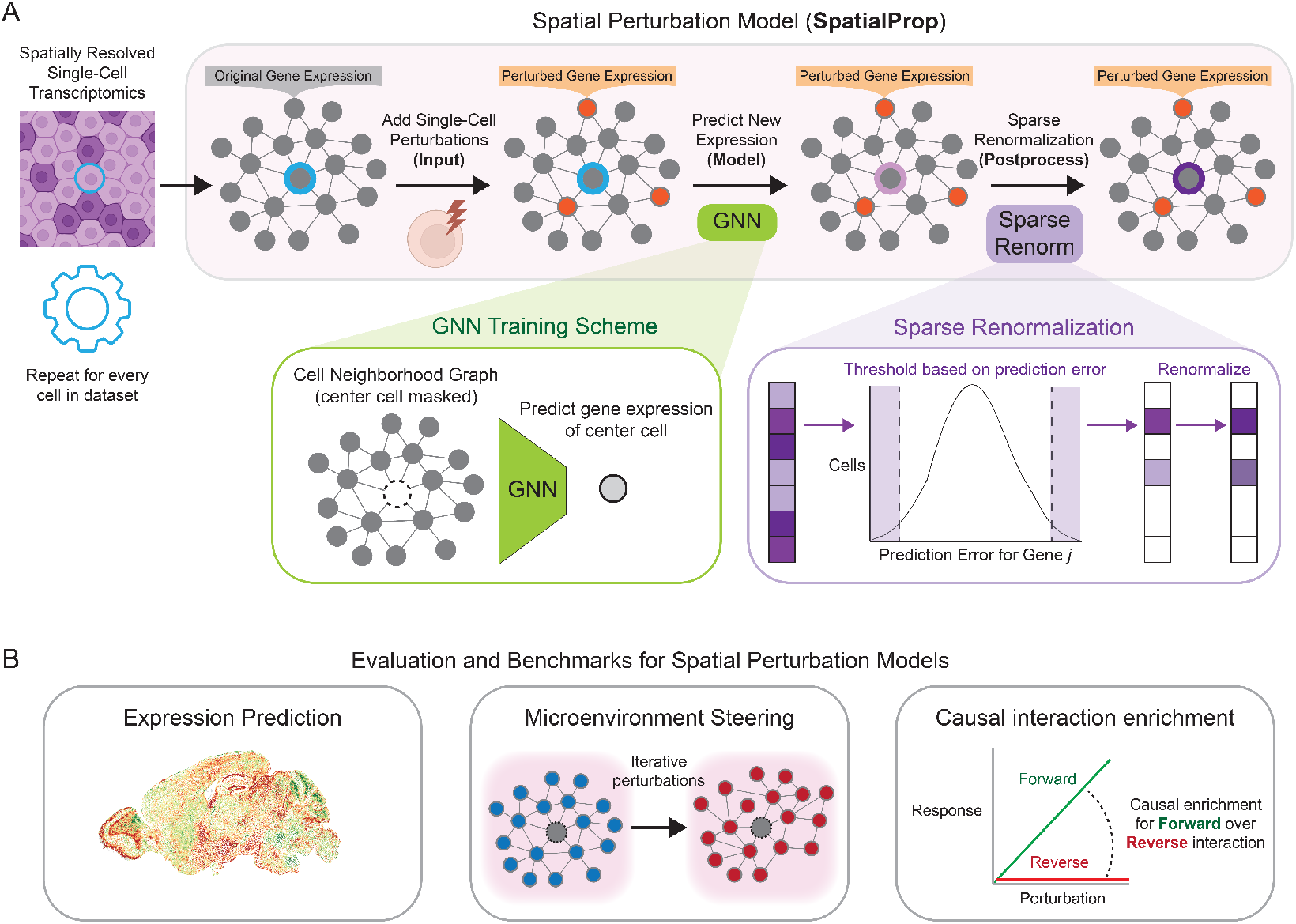
SpatialProp pipeline and evaluation. (A) Schematic illustration of the SpatialProp pipeline, which receives a spatial transcriptomics dataset and a set of single-cell perturbations as input and then generates the predicted perturbed gene expression for every cell based on its tissue microenvironment (represented as a graph). (B) High-level overview of the main evaluation frameworks for assessing performance of spatial perturbation models like SpatialProp.

### Graph neural network accurately predicts single-cell gene expression from tissue microenvironments

The core module of SpatialProp is a graph neural network (GNN) that is trained to predict the gene expression of a cell from the gene expression of neighboring cells in its tissue microenvironment with an intermediate step of predicting center cell type (Fig. 1A, see Methods for further details). The model is trained on an independent spatially resolved single-cell transcriptomics dataset, which permits lightweight and context-specific modeling (see Table 1 and Table 2 for datasets). To extract tissue microenvironments, we devised a sampling strategy to ensure fair representation of all annotated cell types in the dataset (see Methods), and constructed 2-hop nearest neighborhood graphs centered around the “center cell.” In each tissue microenvironment, cells are represented as nodes, with gene expression encoded within node features, and spatial proximity is represented by edges between nodes.

**Table 1:** Overview of spatial transcriptomics datasets.

| Dataset name | Identifier | Reference | Species | Tissue | Spatial technology | Number profiled genes | Number profiled cells |
| --- | --- | --- | --- | --- | --- | --- | --- |
| MERFISH-AgingCoronal | AC | Sun et al., 2025 [30] | *Mus musculus* | Brain | MERFISH | 300 | 1453144 |
| MERFISH-AgingSagittal | AS | Sun et al., 2025 [30] | *Mus musculus* | Brain | MERFISH | 300 | 860717 |
| MERFISH-LocalInjury | MS | Androvic et al., 2023 [31] | *Mus musculus* | Brain | MERFISH | 287 | 280176 |
| MERFISH-Exercise | EX | Sun et al., 2025 [30] | *Mus musculus* | Brain | MERFISH | 300 | 901905 |
| ISS-GlobalInjury | EA | Kukanja et al., 2024 [32] | *Mus musculus* | CNS | ISS | 239 | 321543 |
| MERFISH-AgingPilot | AP | Sun et al., 2024 [27] | *Mus musculus* | Brain | MERFISH | 140 | 198867 |
| MERFISH-Reprogramming | RE | Sun et al., 2025 [30] | *Mus musculus* | Brain | MERFISH | 300 | 1015177 |
| STARmap-Alzheimers | AD | Zeng et al., 2023 [33] | *Mus musculus* | Brain | STARmap PLUS | 2766 | 72165 |
| MERFISH-HeartDevelopment | HD | Farah et al., 2024 [34] | *Homo sapiens* | Heart | MERFISH | 238 | 152720 |

**Table 2:**
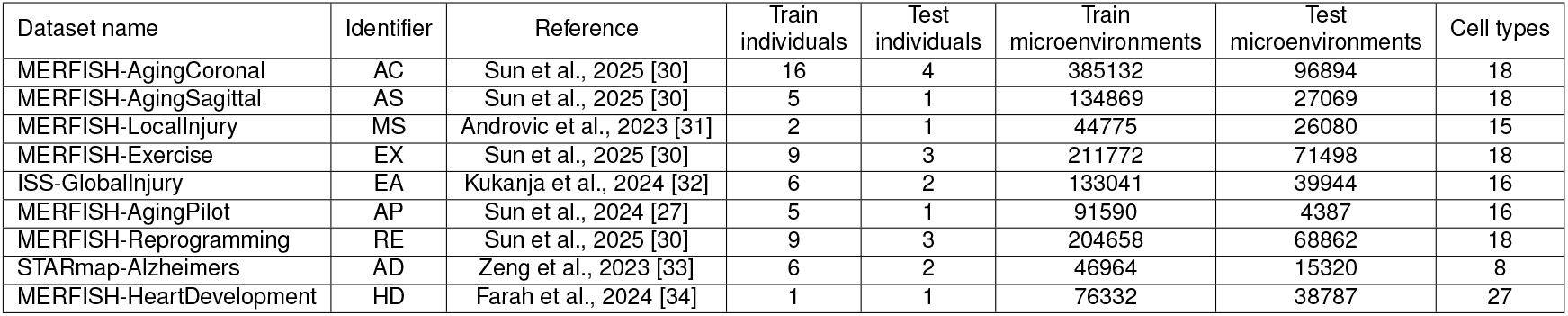
Overview of training and evaluation datasets for each SpatialProp model.

| Dataset name | Identifier | Reference | Train individuals | Test individuals | Train microenvironments | Test microenvironments | Cell types |
| --- | --- | --- | --- | --- | --- | --- | --- |
| MERFISH-AgingCoronal | AC | Sun et al., 2025 [30] | 16 | 4 | 385132 | 96894 | 18 |
| MERFISH-AgingSagittal | AS | Sun et al., 2025 [30] | 5 | 1 | 134869 | 27069 | 18 |
| MERFISH-LocalInjury | MS | Androvic et al., 2023 [31] | 2 | 1 | 44775 | 26080 | 15 |
| MERFISH-Exercise | EX | Sun et al., 2025 [30] | 9 | 3 | 211772 | 71498 | 18 |
| ISS-GlobalInjury | EA | Kukanja et al., 2024 [32] | 6 | 2 | 133041 | 39944 | 16 |
| MERFISH-AgingPilot | AP | Sun et al., 2024 [27] | 5 | 1 | 91590 | 4387 | 16 |
| MERFISH-Reprogramming | RE | Sun et al., 2025 [30] | 9 | 3 | 204658 | 68862 | 18 |
| STARmap-Alzheimers | AD | Zeng et al., 2023 [33] | 6 | 2 | 46964 | 15320 | 8 |
| MERFISH-HeartDevelopment | HD | Farah et al., 2024 [34] | 1 | 1 | 76332 | 38787 | 27 |

We independently trained the SpatialProp GNN on a variety of spatially resolved single-cell transcriptomics datasets detailed in Table 1 and evaluated its predictive performance on held-out test sets consisting of whole sections from biological replicates. Generally, we observed strong concordance between the predicted gene expression and the ground truth gene expression of the center cell in the test set across multiple independent datasets, with the SpatialProp GNN generally outperforming a mean gene expression baseline computed over the training examples (Fig. 2A). We also note similar performance compared to a strong *k*-hop mean baseline, which is the mean gene expression across cells in the corresponding tissue microenvironment of the test set, and to several more complex GNN model architectures (Appendix A). The performance of the model is similar across different cell types (Fig. 2B), suggesting generalization to rare cell types in a tissue. The complete SpatialProp pipeline can flexibly model perturbations and predict tissue-wide perturbation responses, as demonstrated for inflammatory signaling in the mouse brain (Fig. 2C, see Appendix B for perturbation and response genes).

**Figure 2:**
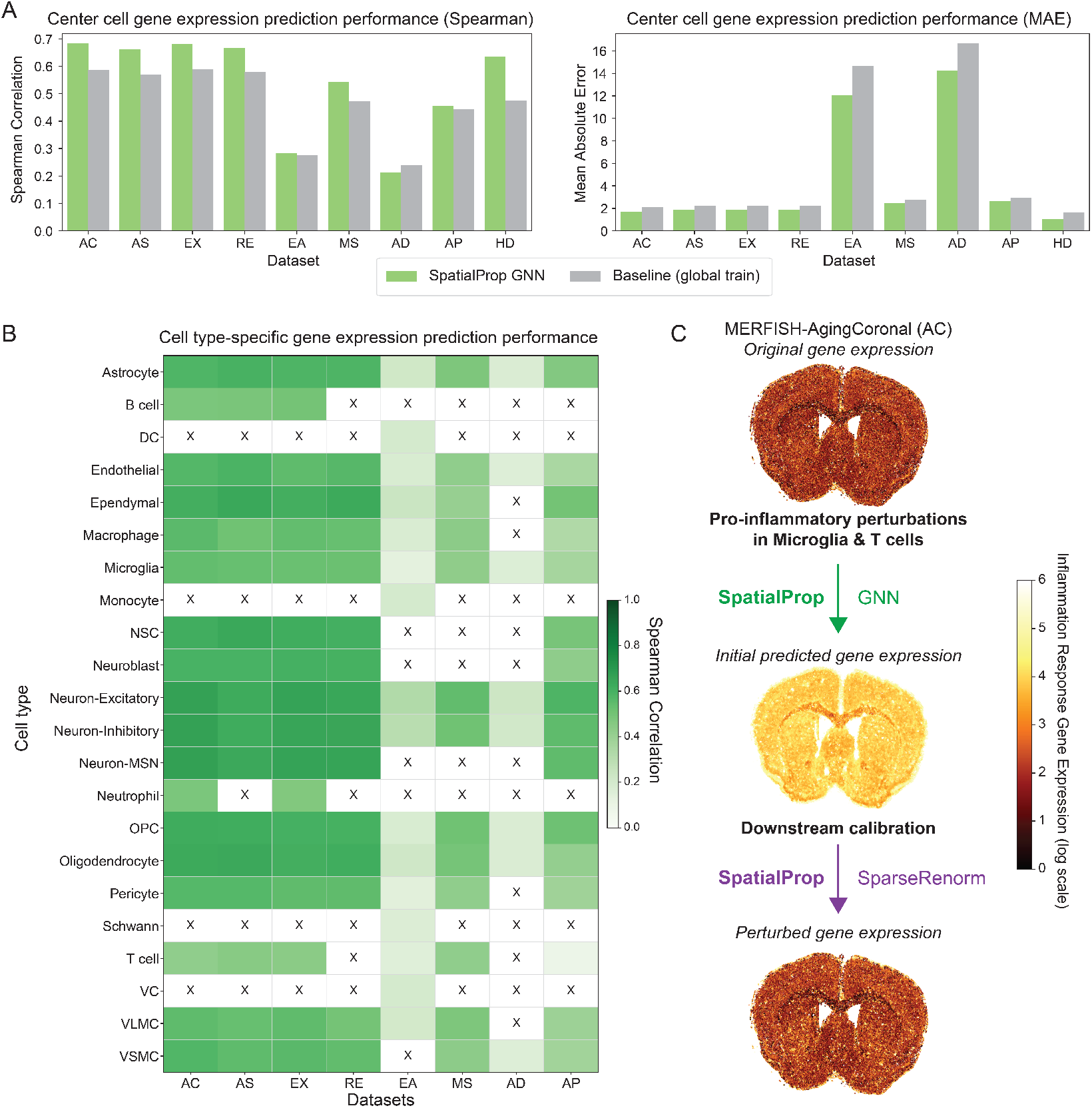
SpatialProp GNN predicts spatial gene expression. (A) Spearman correlation (left) and mean absolute error (right) between predicted gene expression and the ground truth gene expression (micro-averaged across cell types) for multiple spatially resolved single-cell transcriptomics datasets. Shown are results from the SpatialProp GNN module and a baseline consisting of global mean gene expression in training data. (B) Heatmap showing cell type-specific Spearman correlation between predicted gene expression from the GNN module and the ground truth gene expression in mouse brain datasets. (C) Spatial visualization of inflammation response gene expression in all cells from a mouse brain section before and after pro-inflammatory perturbations in microglia and T cells and using SpatialProp for predictions. List of perturbation and response genes provided in Appendix B.

### SpatialProp maps fine-grained steering of tissue microenvironments towards target states

Given that deep learning methods for predicting gene expression from spatial and single-cell transcriptomics data can be prone to fixating on cell type as a surrogate for gene expression [24], we devised a tissue microenvironment steering experiment to test the capacity of SpatialProp to predict the effect of single-cell perturbations to the tissue microenvironment of cells from the same cell type (see Methods). The tissue microenvironment steering experiment setup consists of a starting graph, representing the tissue microenvironment of the starting cell, and a target graph, representing the tissue microenvironment of the target cell. Through successive perturbations consisting of sampling some cells from the target graph to replace cells in the starting graph, the steering experiment transitions the tissue microenvironment of the starting cell to be more similar to that of the target cell (Fig. 3A). Across multiple datasets, SpatialProp recovered increasing similarity between the predicted perturbed expression of the starting cell and the predicted gene expression of the target cell for successively larger steering perturbations to the starting microenvironment (Fig. 3B). These results suggest that SpatialProp reliably predicts the effect of fine-grained perturbations in tissue microenvironments and captures the influence of the tissue microenvironment on center cell gene expression in a manner that is independent of cell type.

**Figure 3:**
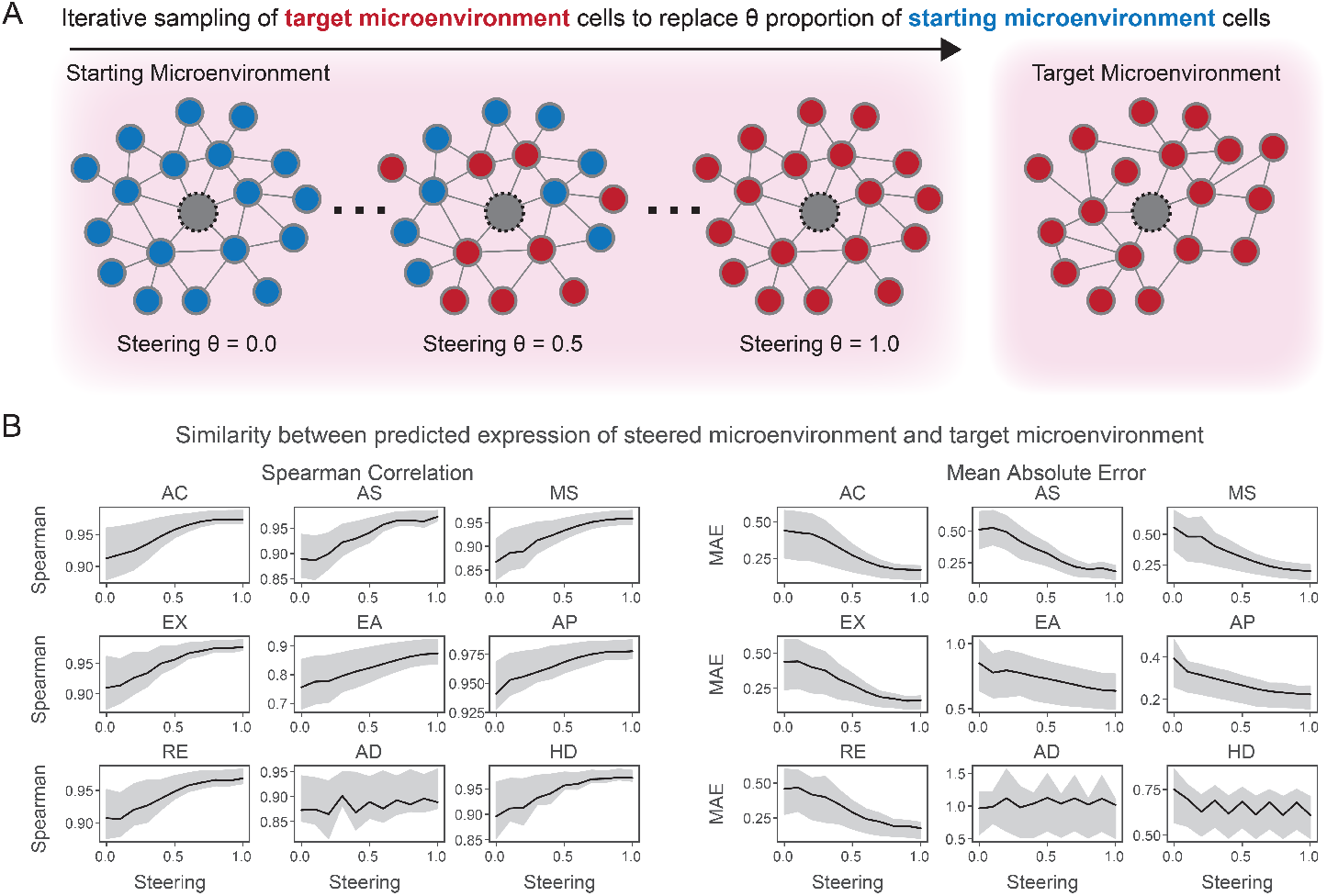
Cell type-specific tissue microenvironment steering experiments. (A) Schematic illustration of the tissue microenvironment steering experiment where given a starting graph and target graph representing tissue microenvironments centered around different cells of the same cell type, the experiment iteratively steers the tissue microenvironment of the starting graph towards that of the target graph by sampling cells from the target tissue microenvironment to replace cells in the starting tissue microenvironment. (B-C) Line plots showing the Spearman correlation (B) or mean absolute error (C) between the SpatialProp-predicted perturbed gene expression of the starting cell compared to the predicted gene expression of the target cell at ten different levels of steering across multiple datasets. Shaded regions for line plots represent 95% confidence interval.

### Evaluating causal enrichment in spatial perturbation predictions

A key question is whether SpatialProp predictions, which are derived from a model trained solely on observational data, are also enriched for causal spatial interactions between cells. Although limited spatial perturbation datasets have become available [18, 19], a systematic benchmarking of known spatial cell-cell interactions is largely absent. To assess causal enrichment in spatial perturbation models, we introduce a new benchmark dataset referred to as CausalInteractionBench. To build CausalInteractionBench, we leverage domain knowledge encoded in the Gene Ontology to extract sets of genes that are annotated for biological processes in the sender (e.g. “production of X”, “X secretion”, etc.) or receiver (e.g. “response to X”, “X signaling”, etc.) pathways of an arbitrary mediator X (Fig. 4A). Examples of mediators include cytokines, metabolites, and secreted lipids or fatty acids (see Appendix C for the full list of mediators). Under the assumption that the majority of these directional interactions proceed only in the forward direction (i.e. sender signals to receiver via X) and not in the reverse direction (i.e. receiver signals to sender via X), we can assess the ability of a predictive model in selectively predicting the effects of perturbations in forward and not reverse directions (Fig. 4AB). We define the Causality Enrichment Score (CES) to quantify the enrichment of SpatialProp predictions that have greater alignment with the forward interaction compared to the reverse interaction across CausalInteractionBench (see Methods for full details).

**Figure 4:**
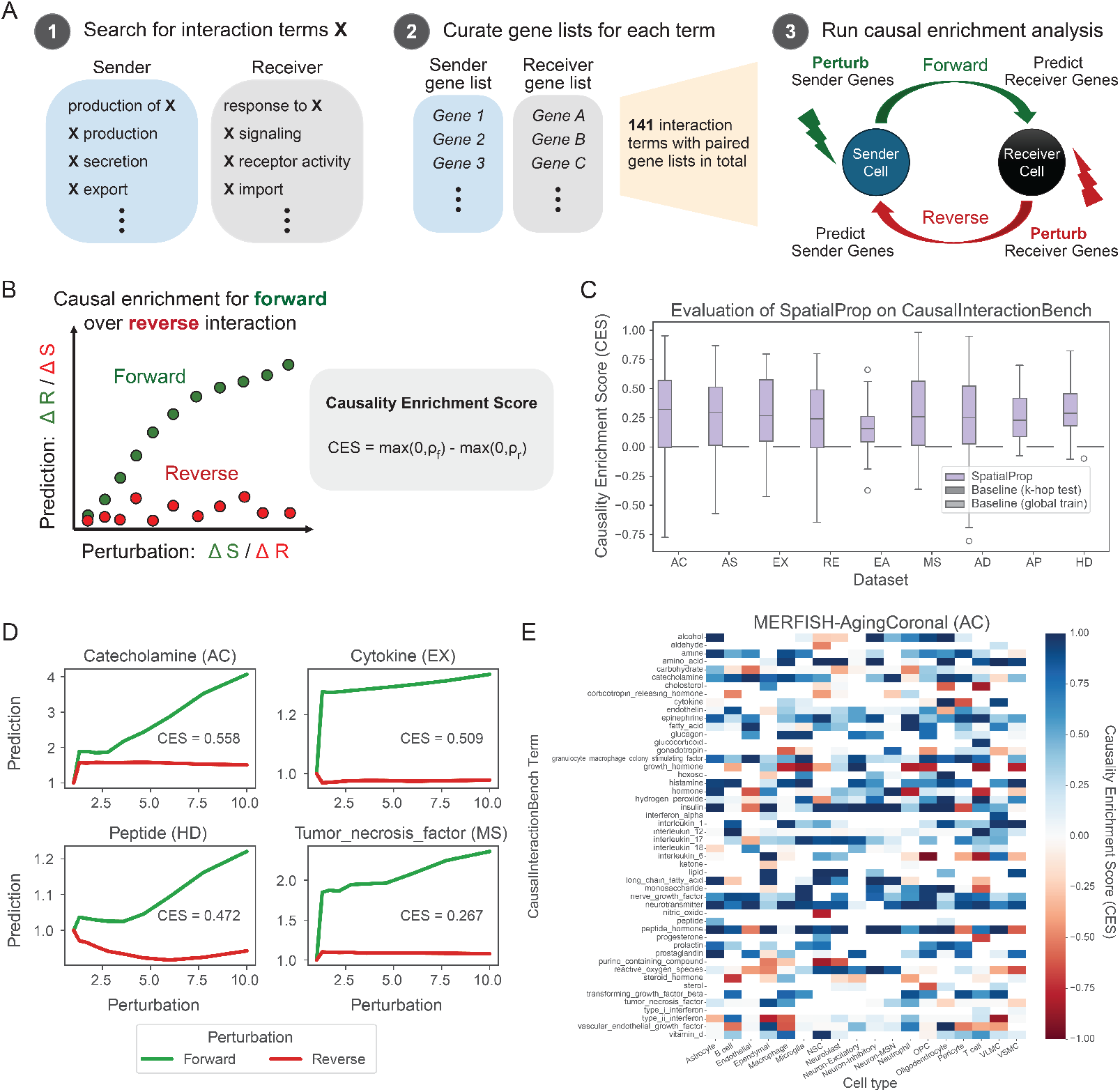
Causal benchmarking for spatial perturbation predictions with CausalInteractionBench. (A) Schematic illustration of the CausalInteractionBench curation process and the evaluation scheme using forward and reverse perturbation of sender and receiver pathways to assess for causal predictions of spatial perturbation effects. (B) Schematic illustration of the response to perturbation curves for a causally enriched interaction and the calculation of the Causality Enrichment Score (CES). (C) Box plots of CES values for SpatialProp and two baselines (global train mean and k-hop test mean) across all CausalInteractionBench terms that are present in the corresponding datasets. Median and interquartile range are delineated in the box and whiskers correspond to 1.5 times the interquartile range. (D) Examples of causally-enriched interaction terms across four datasets. Red line indicates the response to forward perturbations and green line indicates the response to reverse perturbations. (E) Heatmap showing Causality Enrichment Score (CES) by CausalInteractionBench term and cell type in the Aging-Coronal dataset.

In total, we compiled 133 sender-receiver interactions in CausalInteractionBench, encompassing a broad range of different interactions including metabolic signaling and secreted factors (Fig. 4A). We evaluated the causal enrichment of SpatialProp predictions for the subset of interactions that are adequately captured in each dataset (at least one gene in the sender gene set and at least one gene in the receiver gene set). Generally, forward responses were more positively correlated than reverse responses with respect to the level of perturbation, with a mean difference Spearman correlation of 0.094. When considering only the interactions with positive correlation between forward perturbation and response (i.e. that the model captures the association between sender and receiver gene lists), we observed generally positive CES for SpatialProp across all evaluation datasets and a large majority of individual interactions, while neutral CES was observed in both the global train mean and k-hop test mean baselines (Fig. 4C). Some representative examples of mediators revealing causal enrichment are shown in Fig. 4D and are supported by prior studies. For example, cholesterol production in astrocytes supports the activity of neurons along with other lipid-related cell-cell interactions in the mouse brain [25]. Some null examples can be found in Appendix D. Interestingly, the CES scores were generally consistent across cell types for a given interaction term (Fig. 4E). Together, these results suggest that the SpatialProp computational pipeline can predict the spatial effects of single-cell perturbations with causal enrichment for known biological interactions.

## Discussion

We introduce SpatialProp as a simple spatial perturbation model that leverages a graph neural network framework to propagate the transcriptomic effects of known single-cell perturbations across an entire tissue section. Compared to previous applications of graph neural networks to predicting spatial gene expression [24], SpatialProp uses a minimal feature set consisting of only gene expression. As a result, SpatialProp models gene expression conditioned solely on tissue microenvironment variability, which improves flexible modeling of perturbation effects. We further develop a set of evaluation frameworks to guide the development of future spatial perturbation models. SpatialProp accurately predicts spatial gene expression, finely maps the transitions between cells of the same cell type via perturbations in the tissue microenvironment, and provides causally-enriched predictions of spatial perturbation effects.

As part of the evaluation framework for SpatialProp, we introduce CausalInteractionBench to assess for causally enriched spatial perturbation predictions. Currently, CausalInteractionBench is limited to general cell-cell interactions inferred from the Gene Ontology. Inclusion of additional interaction gene lists through autonomous procurement from existing literature or other databases can serve to broaden the representation of different cell-cell interactions in CausalInteractionBench and also extend our benchmarking to cell type-specific interactions. CausalInteractionBench may be further complemented with readouts from spatial perturbation assays such as spatial Perturb-seq.

SpatialProp is synergistic with recent efforts towards predicting single-cell perturbation effects [8, 10, 11, 26]. SpatialProp takes perturbed single-cell transcriptomes as input, which can either be from experimental readouts such as Perturb-seq or from the predictions of single-cell perturbation models. As such, improvements in predicting the effects of perturbations on single-cell gene expression will also expand the utility of SpatialProp. When deployed in conjunction with existing single-cell perturbation models, Spatial-Prop provides an end-to-end pipeline for *in silico* simulation of tissue-wide gene expression in response to precise single-cell gene expression perturbations.

Currently, SpatialProp is independently trained on each spatial transcriptomics dataset to enable prediction of spatial perturbation effects. Training SpatialProp across a large-scale, integrated spatial transcriptomics atlas will be necessary to build a universal spatial perturbation model that can be deployed across different contexts without retraining. However, the integration of spatial transcriptomics datasets, particularly those generated from imaging-based approaches, is difficult given the low overlap between different gene panels. In future settings, this limitation may be overcome by using spatial transcriptomics imputation [27] or similar approaches [28] to permit integration of datasets across different species, organs, and tissues and unlock the universal prediction of spatial perturbation effects.

Finally, SpatialProp is limited in its training on observational spatial transcriptomics, and future iterations that leverage data from new developments in spatial perturbation assays are likely to exhibit improved performance in predicting the spatial effects of single-cell perturbations. In that regard, the SpatialProp framework and associated benchmarks may guide further developments in this space.

## Methods

### Spatial transcriptomics training and evaluation datasets

We independently trained and evaluated SpatialProp on datasets spanning multiple tissue types and spatial technologies, detailed in Table 1. To ensure balanced representation of center cell types in the tissue microenvironments across training and test datasets, we performed a two-step sampling procedure. First, we sampled 100 cells per cell type to serve as the first set of center cells, retaining all cell types. We then constructed a Delaunay triangulation graph across each sample using Squidpy [29], pruning long-range edges greater than 200 µm. In the graph, each node is a cell and has features corresponding to its normalized gene expression such that the total expression per cell is equal to the number of genes. We induced the 2-hop neighborhood graphs centered at each of these center cells, in which each graph represents a tissue microenvironment. In the second step, we further induced 2-hop neighborhood graphs around each of the surrounding cells in these tissue microenvironments and added these graphs as additional tissue microenvironments in the training and test datasets. The first step ensured balanced representation of rare cell types, while the second step augmented training and test dataset sizes. Individual samples were assigned randomly for training and testing splits during model development. The number of individuals and tissue microenvironments is reported for the training and evaluation datasets in Table 2.

### Spatial propagation of single-cell perturbations (SpatialProp)

The SpatialProp framework takes as input a spatially resolved single-cell transcriptomics dataset consisting of the gene expression measurements and spatial coordinates of each cell. Then, for each cell, SpatialProp extracts its induced 2-hop neighborhood to build a tissue microenvironment graph and masks the node features corresponding to the center cell. Using the underlying GNN model (see subsection below for details), a prediction of the gene expression for the center cell is made using the tissue microenvironment graph (prediction referred to as “base prediction”). Perturbations are then made to the tissue microenvironment, which can be user-defined and consist of any number of gene expression perturbations. The underlying GNN model is used to predict the gene expression of the center cell using the perturbed tissue microenvironment graph (prediction referred to as “initial prediction”). These predictions and perturbations are then performed across all cells in the dataset.

To guard against model prediction errors, we make the assumption that most spatial perturbation effects can be represented by sparse changes in gene expression and post-process the initial predictions accordingly using a two-step approach that we refer to as SparseRenorm. First, we construct a sparse mask to update the original gene expression with GNN-predicted gene expression. For each gene, an empirical distribution of prediction errors is constructed by subtracting the base prediction from the ground truth gene expression. Then, a final prediction of perturbed gene expression is constructed for each cell, where the initial prediction is used if the difference between the initial prediction is more extreme than the prediction error for that cell and is also more extreme than the bottom 5% or top 5% of prediction errors in the empirical distribution. Otherwise, the ground truth expression is retained for that cell. This procedure is repeated independently for each gene. As a result, this first step provides an aggressive sparse updating of the ground truth gene expression to account for perturbation. In the second step of SparseRenorm, the output of the first step is renormalized by multiplying all expression values in a cell by a constant such that total gene expression is conserved between the original (unperturbed) gene expression and the output of the second step for that cell. The final output after both steps is returned as the predicted perturbed expression generated by the full SpatialProp pipeline.

### Graph deep learning prediction of gene expression from tissue microenvironment

The predictive module used in SpatialProp consists of a GNN model trained to predict center cell gene expression from the gene expression of surrounding cells in the tissue microenvironment. The GNN consists of convolutional layers of a graph isomorphism network (GIN) [35], implemented with PyTorch Geometric. Network depth is set equal to *k*, which defines the *k*-hop neighborhood for tissue microenvironments induced around center cells. In all SpatialProp settings presented in this study, we set *k* = 2 with the exception of *k* = 3 used in additional benchmarking. The final prediction is made using the embedding of the center cell, which is masked (i.e. gene expression set to zero) at the start of training. Parameters of the GNN are trained with Adam optimizer, learning rate of 10^*−*4^, and a weighted mean absolute error loss to adjust for zero inflation by specifying different weighting of (*w*_0_, *w*_1_)for zero and non-zero ground truth gene expression values respectively:

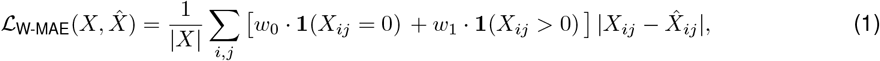

where *X* is the ground truth cell by gene expression matrix of the center cells across a batch and 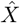 is the GNN-predicted center cell gene expression. We set *w*_0_ = 1 and *w*_1_ = 10. Each SpatialProp model is trained over 50 epochs with no early stopping. The model after 50 epochs is evaluated on held-out tissue microenvironments in predicting masked center cell gene expression. We evaluated performance by computing Spearman correlation and mean absolute error with respect to the ground truth center cell gene expression.

We also explored and benchmarked more complex GNN architectures including those with global attention or sum pooling strategies for graph-level prediction. These further included both multi-task and nested architectures to predict surrogate targets such as cell type in addition to gene expression. Generally, these more complex architectures did not outperform the simple GNN model for the prediction task (Appendix A).

### Tissue microenvironment steering setup

The goal of the tissue microenvironment steering experiment is to assess whether a spatial perturbation model like SpatialProp can adequately model intra-cell type heterogeneity in gene expression conditioned on the tissue microenvironment. The setup consists of successive perturbations that steer a starting graph, representing the tissue microenvironment of the starting cell, towards a target graph, representing the tissue microenvironment of the target cell, and assessing the change in similarity between the perturbed expression of the steered graph and the target graph as predicted by SpatialProp. In the steering experiment, perturbations are defined by the random replacement of cells in the starting graph with randomly sampled cells from the target graph. The perturbation level *θ* is defined as the proportion of cells in the starting graph that are replaced. For higher perturbation levels, the distribution of cells in the starting graph will resemble that of the target graph. To evaluate for similarity in the predicted perturbed expression, we compute the Spearman correlation and the mean absolute error between the perturbed expression predicted for the steered starting graph and the expression predicted for the target graph by SpatialProp. Only renormalization is performed in the SparseRenorm step because no perturbation was performed for target graph expression prediction. For evaluation, we assess whether predictions from SpatialProp, or another spatial perturbation model, increases in similarity as the steering level *θ* increases (i.e. *θ →* 1).

### CausalInteractionBench: benchmark for evaluation of causal spatial perturbation predictions

To build CausalInteractionBench, we constructed gene sets encoding known spatial cell-cell interactions to evaluate the causal enrichment of SpatialProp predictions. Each interaction is defined by a signal produced by the “sender” cell that elicits a response from the “receiver” cell, with corresponding gene sets for the sender and receiver. We autonomously discovered key interaction terms by performing a string matching search across all Gene Ontology Biological Process (GO BP) terms. For an arbitrary interaction term (i.e. mediator) denoted *X*, we included all genes in GO BP terms corresponding to production, synthesis, secretion, and export of *X* to construct the sender gene set. We included all genes in GO BP terms corresponding to response, activity, signaling, and receptor binding of *X* to construct the receiver gene set. This process was iterated for all autonomously discovered *X* terms with at least one GO BP term in the sender gene set and one GO BP term in the receiver gene set. We accessed the Gene Ontology through the latest versions of go-basic.obo and mgi.gaf as of July 21, 2025 through https://geneontology.org/. A table of interaction terms and GO identifiers used in CausalInteractionBench can be found in Appendix C. This table and the complete gene lists for each interaction are also available at https://github.com/sunericd/CausalInteractionBench.

CausalInteractionBench can be used to evaluate spatial perturbation models like SpatialProp by measuring perturbation responses in the forward (sender-to-receiver) and reverse (receiver-to-sender) directions and comparing the relative enrichment for the forward perturbation response compared to reverse perturbation response. The intuition for this bidirectional comparison of perturbation responses is that the forward direction corresponds to a causal interaction while there is no expectation that the reverse interaction would exist in the real world. To generate perturbations in the tissue microenvironment, we scale the expression of all genes in either the sender gene set (forward direction) or the receiver gene set (reverse direction) by a perturbation multiplier (referred to as “perturbation level”) ranging from 1 to 10 in all cells except for the center cell. Then, we measure the perturbed gene expression as predicted by SpatialProp for the center cell after the perturbation in the tissue microenvironment and compute a normalized response with respect to the predicted response under non-perturbed conditions. This approach is iterated for different perturbation levels and aggregated across all tissue microenvironments to compute a Causality Enrichment Score (CES) for a given interaction (see following section for details). For each dataset, this setup was used to evaluate SpatialProp on all interactions with at least one gene represented in the sender gene set and one gene represented in the receiver gene set.

### Causality enrichment score

To quantify the causal enrichment of SpatialProp for the forward interaction compared to the reverse interaction, we define the Causality Enrichment Score (CES):

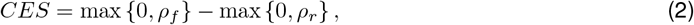

where *ρ*_*f*_ is the Spearman correlation between the perturbation level and the normalized forward response, and *ρ*_*r*_ is the Spearman correlation between the perturbation level and the normalized reverse response. The normalized forward response is calculated as the predicted forward response (i.e. sum of predicted gene expression in the receiver gene set of the center cell after perturbation in the sender gene set in all other cells of the tissue microenvironment) divided by the starting forward response (i.e. sum of gene expression in the receiver gene set of the center cell without perturbation). The normalized reverse response is calculated as the predicted reverse response (i.e. sum of predicted gene expression in the sender gene set of the center cell after perturbation in the receiver gene set in all other cells of the tissue microenvironment) divided by the starting reverse response (i.e. sum of gene expression in the sender gene set of the center cell without perturbation).

## Code availability

The SpatialProp software package for training and deploying the model across user-defined perturbations and associated code for all analyses are available at https://github.com/abuendia/spatial-prop.

## Acknowledgments

We thank Anshul Kundaje for helpful discussions on the model and baselines. Funding support was provided by Knight-Hennessy Scholars program (to E.D. Sun), Paul and Daisy Soros Fellowship for New Americans (to E.D. Sun), the National Science Foundation Graduate Research Fellowship Program (to E.D. Sun), NIH training grant T15LM007033 (to A. Buendia), NIH grant R01AG071711 (to A. Brunet), the Milky Way Research Foundation (to A. Brunet), Simons Foundation grant (to A. Brunet), the Knight Initiative for Brain Resilience (to A. Brunet), a generous gift from M. and T. Barakett (to A. Brunet), CZI collaborative pairs pilot project award (to A. Brunet and J. Zou), Chan Zuckerberg Biohub–San Francisco Investigator program (to A. Brunet and J. Zou), NSF CAREER 1942926 (to J. Zou), and grants from Silicon Valley Foundation (to J. Zou).

## References

[1] Atray Dixit et al. “Perturb-Seq: Dissecting Molecular Circuits with Scalable Single-Cell RNA Profiling of Pooled Genetic Screens”. en. In: Cell 167.7 (Dec. 2016), 1853–1866.e17. ISSN: 00928674. DOI: 10.1016/j.cell.2016.11.038. URL: https://linkinghub.elsevier.com/retrieve/pii/S0092867416316105.

[2] Britt Adamson et al. “A Multiplexed Single-Cell CRISPR Screening Platform Enables Systematic Dissection of the Unfolded Protein Response”. en. In: Cell 167.7 (Dec. 2016), 1867–1882.e21. ISSN: 00928674. DOI: 10.1016/j.cell.2016.11.048. URL: https://linkinghub.elsevier.com/retrieve/pii/S0092867416316609.

[3] Paul Datlinger et al. “Pooled CRISPR screening with single-cell transcriptome readout”. en. In: Nature Methods 14.3 (Mar. 2017), pp. 297–301. ISSN: 1548-7091, 1548-7105. DOI: 10.1038/nmeth.4177. URL: https://www.nature.com/articles/nmeth.4177.

[4] Diego Adhemar Jaitin et al. “Dissecting Immune Circuits by Linking CRISPR-Pooled Screens with Single-Cell RNA-Seq”. en. In: Cell 167.7 (Dec. 2016), 1883–1896.e15. ISSN: 00928674. DOI: 10.1016/j.cell.2016.11.039. URL: https://linkinghub.elsevier.com/retrieve/pii/S0092867416316117.

[5] Joseph M. Replogle et al. “Mapping information-rich genotype-phenotype landscapes with genomescale Perturb-seq”. en. In: Cell 185.14 (July 2022), 2559–2575.e28. ISSN: 00928674. DOI: 10.1016/j.cell.2022.05.013. URL: https://linkinghub.elsevier.com/retrieve/pii/S0092867422005979.

[6] Thomas M. Norman et al. “Exploring genetic interaction manifolds constructed from rich single-cell phenotypes”. en. In: Science 365.6455 (Aug. 2019), pp. 786–793. ISSN: 0036-8075, 1095-9203. DOI: 10.1126/science.aax4438. URL: https://www.science.org/doi/10.1126/science.aax4438.

[7] Mohammad Lotfollahi, F. Alexander Wolf, and Fabian J. Theis. “scGen predicts single-cell perturbation responses”. en. In: Nature Methods 16.8 (Aug. 2019), pp. 715–721. ISSN: 1548-7091, 1548-7105. DOI: 10.1038/s41592-019-0494-8. URL: https://www.nature.com/articles/s41592-019-0494-8.

[8] Yusuf Roohani, Kexin Huang, and Jure Leskovec. “Predicting transcriptional outcomes of novel multigene perturbations with GEARS”. en. In: Nature Biotechnology 42.6 (June 2024), pp. 927–935. ISSN: 1087-0156, 1546-1696. DOI: 10.1038/s41587-023-01905-6. URL: https://www.nature.com/articles/s41587-023-01905-6.

[9] Charlotte Bunne et al. “Learning single-cell perturbation responses using neural optimal transport”. en. In: Nature Methods 20.11 (Nov. 2023), pp. 1759–1768. ISSN: 1548-7091, 1548-7105. DOI: 10.1038/s41592-023-01969-x. URL: https://www.nature.com/articles/s41592-023-01969-x.

[10] Haotian Cui et al. “scGPT: toward building a foundation model for single-cell multi-omics using generative AI”. en. In: Nature Methods 21.8 (Aug. 2024), pp. 1470–1480. ISSN: 1548-7091, 1548-7105. DOI: 10.1038/s41592-024-02201-0. URL: https://www.nature.com/articles/s41592-024-02201-0.

[11] Minsheng Hao et al. “Large-scale foundation model on single-cell transcriptomics”. en. In: Nature Methods 21.8 (Aug. 2024), pp. 1481–1491. ISSN: 1548-7091, 1548-7105. DOI: 10.1038/s41592-024-02305-7. URL: https://www.nature.com/articles/s41592-024-02305-7.

[12] Constantin Ahlmann-Eltze, Wolfgang Huber, and Simon Anders. “Deep-learning-based gene perturbation effect prediction does not yet outperform simple linear baselines”. en. In: Nature Methods 22.8 (Aug. 2025), pp. 1657–1661. ISSN: 1548-7091, 1548-7105. DOI: 10.1038/s41592-025-02772-6. URL: https://www.nature.com/articles/s41592-025-02772-6.

[13] Daniel R Wong, Abby S Hill, and Rob Moccia. “Simple controls exceed best deep learning algorithms and reveal foundation model effectiveness for predicting genetic perturbations”. en. In: Bioinformatics 41.6 (June 2025). Ed. by Anthony Mathelier, btaf317. ISSN: 1367-4811. DOI: 10.1093/bioinformatics/btaf317. URL: https://academic.oup.com/bioinformatics/article/doi/10.1093/bioinformatics/btaf317/8142305.

[14] Gabriel M. Mejia et al. Diversity by Design: Addressing Mode Collapse Improves scRNA-seq Perturbation Modeling on Well-Calibrated Metrics. arXiv:2506.22641 [q-bio]. June 2025. DOI: 10.48550/arXiv.2506.22641. URL: http://arxiv.org/abs/2506.22641.

[15] Euxhen Hasanaj et al. Multimodal Benchmarking of Foundation Model Representations for Cellular Perturbation Response Prediction. en. June 2025. DOI: 10.1101/2025.06.26.661186. URL: http://biorxiv.org/lookup/doi/10.1101/2025.06.26.661186.

[16] Yan Wu et al. PerturBench: Benchmarking Machine Learning Models for Cellular Perturbation Analysis. arXiv:2408.10609 [cs]. June 2025. DOI: 10.48550/arXiv.2408.10609. URL: http://arxiv.org/abs/2408.10609.

[17] Ihab Bendidi et al. Benchmarking Transcriptomics Foundation Models for Perturbation Analysis : one PCA still rules them all. arXiv:2410.13956 [cs]. Nov. 2024. DOI: 10.48550/arXiv.2410.13956. URL: http://arxiv.org/abs/2410.13956.

[18] LoÏc Binan et al. “Simultaneous CRISPR screening and spatial transcriptomics reveal intracellular, intercellular, and functional transcriptional circuits”. en. In: Cell 188.8 (Apr. 2025), 2141–2158.e18. ISSN: 00928674. DOI: 10.1016/j.cell.2025.02.012. URL: https://linkinghub.elsevier.com/retrieve/pii/S0092867425001977.

[19] Reuben A. Saunders et al. “Perturb-Multimodal: A platform for pooled genetic screens with imaging and sequencing in intact mammalian tissue”. en. In: Cell 188.17 (Aug. 2025), 4790–4809.e22. ISSN: 00928674. DOI: 10.1016/j.cell.2025.05.022. URL: https://linkinghub.elsevier.com/retrieve/pii/S0092867425005720.

[20] Alev Baysoy et al. Spatially Resolved in vivo CRISPR Screen Sequencing via Perturb-DBiT. en. Nov. 2024. DOI: 10.1101/2024.11.18.624106. URL: http://biorxiv.org/lookup/doi/10.1101/2024.11.18.624106.

[21] Zitong Jerry Wang et al. “Identifying perturbations that boost T-cell infiltration into tumours via counterfactual learning of their spatial proteomic profiles”. en. In: Nature Biomedical Engineering 9.3 (Mar. 2025), pp. 390–404. ISSN: 2157-846X. DOI: 10.1038/s41551-025-01357-0. URL: https://www.nature.com/articles/s41551-025-01357-0.

[22] Yuning You et al. Building Foundation Models to Characterize Cellular Interactions via Geometric Self-Supervised Learning on Spatial Genomics. en. Jan. 2025. DOI: 10.1101/2025.01.25.634867. URL: http://biorxiv.org/lookup/doi/10.1101/2025.01.25.634867.

[23] Amir Akbarnejad et al. Mapping and reprogramming human tissue microenvironments with MintFlow. en. June 2025. DOI: 10.1101/2025.06.24.661094. URL: http://biorxiv.org/lookup/doi/10.1101/2025.06.24.661094.

[24] David S. Fischer, Anna C. Schaar, and Fabian J. Theis. “Modeling intercellular communication in tissues using spatial graphs of cells”. en. In: Nature Biotechnology 41.3 (Mar. 2023), pp. 332–336. ISSN: 1087-0156, 1546-1696. DOI: 10.1038/s41587-022-01467-z. URL: https://www.nature.com/articles/s41587-022-01467-z.

[25] Eric D. Sun et al. “Brain aging and rejuvenation at single-cell resolution”. eng. In: Neuron 113.1 (Jan. 2025), pp. 82–108. ISSN: 1097-4199. DOI: 10.1016/j.neuron.2024.12.007.

[26] Abhinav K. Adduri et al. Predicting cellular responses to perturbation across diverse contexts with State. en. June 2025. DOI: 10.1101/2025.06.26.661135. URL: http://biorxiv.org/lookup/doi/10.1101/2025.06.26.661135.

[27] Eric D. Sun et al. “TISSUE: uncertainty-calibrated prediction of single-cell spatial transcriptomics improves downstream analyses”. en. In: Nature Methods 21.3 (Mar. 2024), pp. 444–454. ISSN: 1548-7091, 1548-7105. DOI: 10.1038/s41592-024-02184-y. URL: https://www.nature.com/articles/s41592-024-02184-y.

[28] Tommaso Biancalani et al. “Deep learning and alignment of spatially resolved single-cell transcriptomes with Tangram”. en. In: Nature Methods 18.11 (Nov. 2021), pp. 1352–1362. ISSN: 1548-7091, 1548-7105. DOI: 10.1038/s41592-021-01264-7. URL: https://www.nature.com/articles/s41592-021-01264-7.

[29] Giovanni Palla et al. “Squidpy: a scalable framework for spatial omics analysis”. en. In: Nature Methods 19.2 (Feb. 2022), pp. 171–178. ISSN: 1548-7091, 1548-7105. DOI: 10.1038/s41592-021-01358-2. URL: https://www.nature.com/articles/s41592-021-01358-2.

[30] Eric D. Sun et al. “Spatial transcriptomic clocks reveal cell proximity effects in brain ageing”. en. In: Nature 638.8049 (Feb. 2025), pp. 160–171. ISSN: 0028-0836, 1476-4687. DOI: 10.1038/s41586-024-08334-8. URL: https://www.nature.com/articles/s41586-024-08334-8.

[31] Peter Androvic et al. “Spatial Transcriptomics-correlated Electron Microscopy maps transcriptional and ultrastructural responses to brain injury”. en. In: Nature Communications 14.1 (July 2023), p. 4115. ISSN: 2041-1723. DOI: 10.1038/s41467-023-39447-9. URL: https://www.nature.com/articles/s41467-023-39447-9.

[32] Petra Kukanja et al. “Cellular architecture of evolving neuroinflammatory lesions and multiple sclerosis pathology”. en. In: Cell 187.8 (Apr. 2024), 1990–2009.e19. ISSN: 00928674. DOI: 10.1016/j.cell.2024.02.030. URL: https://linkinghub.elsevier.com/retrieve/pii/S0092867424002332.

[33] Hu Zeng et al. “Integrative in situ mapping of single-cell transcriptional states and tissue histopathology in a mouse model of Alzheimer’s disease”. en. In: Nature Neuroscience 26.3 (Mar. 2023), pp. 430–446. ISSN: 1097-6256, 1546-1726. DOI: 10.1038/s41593-022-01251-x. URL: https://www.nature.com/articles/s41593-022-01251-x.

[34] Elie N. Farah et al. “Spatially organized cellular communities form the developing human heart”. en. In: Nature 627.8005 (Mar. 2024). Publisher: Nature Publishing Group, pp. 854–864. ISSN: 1476-4687. DOI: 10.1038/s41586-024-07171-z. URL: https://www.nature.com/articles/s41586-024-07171-z (visited on 07/29/2025).

[35] Keyulu Xu et al. How Powerful are Graph Neural Networks? arXiv:1810.00826 [cs]. Feb. 2019. DOI: 10.48550/arXiv.1810.00826. URL: http://arxiv.org/abs/1810.00826.

