## Appendix for "SpatialProp: tissue perturbation modeling with spatially resolved single-cell transcriptomics"

Table of Contents


| Appendix | Description |
|---|---|
| A | Supplementary figures showing comprehensive benchmarking of the SpatialProp GNN module against the strong baseline of k-hop mean gene expression on the test graph and more complex GNN architectures. |
| B | List of genes and perturbation setup for mouse brain inflammatory signaling showcase of SpatialProp pipeline. |
| C | Table of interaction terms in CausalInteractionBench and associated Gene Ontology identifiers |
| D | Supplementary figures showing representative interaction terms with approximately zero or negative causal enrichment scores (CES) |

**Appendix A.** Tables showing comprehensive benchmarking of the SpatialProp GNN module against the train global mean baseline, strong baseline of k-hop mean gene expression on the test graph, and more complex GNN architectures.

Spearman correlation between predicted and ground truth center cell gene expression

| Dataset Identifier | SpatialProp (GNN) | Train Global Mean Baseline | Test 2-hop Mean Baseline | GNN (Nested with cell type prediction) | GNN (Cell type with global attention pool) | GNN (Cell type with predicted residual over 2-hop mean baseline) | GNN (Cell type with 3-hop graphs) |
|---|---|---|---|---|---|---|---|
| AC | 0.684 | 0.586 | 0.660 | 0.682 | 0.680 | 0.631 | 0.680 |
| AS | 0.662 | 0.570 | 0.653 | 0.658 | 0.655 | 0.616 | 0.656 |
| EX | 0.681 | 0.589 | 0.666 | 0.678 | 0.678 | 0.632 | 0.676 |
| RE | 0.667 | 0.578 | 0.650 | 0.664 | 0.662 | 0.613 | 0.663 |
| EA | 0.282 | 0.276 | 0.286 | 0.274 | 0.265 | 0.210 | 0.267 |
| MS | 0.543 | 0.473 | 0.603 | 0.529 | 0.533 | 0.546 | 0.515 |
| AD | 0.212 | 0.240 | 0.203 | 0.227 | 0.213 | 0.093 | 0.220 |
| AP | 0.454 | 0.444 | 0.454 | 0.452 | 0.451 | 0.418 | 0.455 |
| HD | 0.634 | 0.476 | 0.590 | 0.631 | 0.631 | 0.574 | 0.631 |

Mean absolute error between predicted and ground truth center cell gene expression

| Dataset Identifier | SpatialProp (GNN) | Train Global Mean Baseline | Test 2-hop Mean Baseline | GNN (Nested with cell type prediction) | GNN (Cell type with global attention pool) | GNN (Cell type with predicted residual over 2-hop mean baseline) | GNN (Cell type with 3-hop graphs) |
|---|---|---|---|---|---|---|---|
| AC | 1.720 | 2.088 | 1.829 | 1.722 | 1.729 | 1.867 | 1.728 |
| AS | 1.855 | 2.244 | 1.945 | 1.863 | 1.872 | 2.017 | 1.878 |
| EX | 1.889 | 2.251 | 1.996 | 1.898 | 1.905 | 2.035 | 1.906 |
| RE | 1.848 | 2.234 | 1.956 | 1.857 | 1.862 | 2.018 | 1.861 |
| EA | 12.059 | 14.655 | 13.604 | 12.111 | 12.251 | 15.714 | 12.202 |
| MS | 2.446 | 2.772 | 2.272 | 2.495 | 2.485 | 2.407 | 2.533 |
| AD | 14.245 | 16.681 | 16.139 | 14.130 | 14.204 | 18.042 | 14.370 |
| AP | 2.650 | 2.936 | 2.846 | 2.660 | 2.649 | 3.039 | 2.610 |
| HD | 1.017 | 1.646 | 1.355 | 1.036 | 1.032 | 1.501 | 1.031 |

**Appendix B.** List of genes and perturbation setup for mouse brain inflammatory signaling showcase of SpatialProp pipeline.

Inflammation Response Genes (response calculated as sum of log1p expression of each gene in the list):

*Stat1, Bst2, Jak1, Ifit1, Cdkn1a, Cdkn2a, C4b, H2-D1, H2-K1*

Pro-inflammatory Perturbation Genes (perturbation modeled as ten-fold increased expression):

*Ifng, Il6, Tnf, Il1a, Il1b, Jun, Apoe, B2m, C1qa, Cd69, Cd9, Lyz2*

**Appendix C.** Table of interaction terms in CausalInteractionBench and associated Gene Ontology identifiers.

| Interaction term | Production GO IDs | Response GO IDs |
|---|---|---|
| maltose | GO:0000024 | GO:0034286, GO:0106081 |
| l-histidine | GO:0000105 | GO:0061460, GO:0090466, GO:1903810 |
| l-tryptophan | GO:0000162, GO:0009096 | GO:1904272 |
| prostaglandin | GO:0001516, GO:0002539, GO:0031392, GO:0031393, GO:0031394, GO:0032306, GO:0032307, GO:0032308, GO:0032310, GO:0071810, GO:0071812 | GO:0034694 |
| histamine | GO:0001693, GO:0001694, GO:0001821, GO:0002349 | GO:0034776, GO:0071420 |
| cytokine | GO:0001816, GO:0001817, GO:0001818, GO:0001819, GO:0002367, GO:0002374, GO:0002375, GO:0002534, GO:0002718, GO:0002719, GO:0002720, GO:0002739, GO:0002740, GO:0002741, GO:0002742, GO:0002743, GO:0002744, GO:0042032, GO:0042035, GO:0042036, | GO:0034097 |

| | | |
|---|---|---|
| | GO:0042089, GO:0042107, GO:0042108, GO:0050663, GO:0050707, GO:0050710, GO:0050715, GO:1900015, GO:1900016, GO:1900017 | |
| serotonin | GO:0001820, GO:0002351, GO:0014062, GO:0014063, GO:0014064, GO:0042427, GO:1905627, GO:1905628, GO:1905629 | GO:1904014, GO:1904015 |
| neurotransmitter | GO:0001956, GO:0007269, GO:0010554, GO:0046928, GO:0046929 | GO:0099601 |
| aldosterone | GO:0002018, GO:0032342, GO:0032347, GO:0032348, GO:0032349, GO:0035932, GO:2000858, GO:2000859, GO:2000860 | GO:1904044, GO:1904045 |
| retinoic acid | GO:0002138, GO:1900052, GO:1900053, GO:1900054 | GO:0032526, GO:0071300 |
| nitric oxide | GO:0002537, GO:0006809, GO:0045019, GO:0045428, GO:0045429 | GO:0071731, GO:0071732 |
| leukotriene | GO:0002540, GO:0019370, GO:0035490, | GO:0061737 |

| | | |
|---|---|---|
| | GO:0035491, GO:0035492 | |
| peptide | GO:0002790, GO:0002791, GO:0002792, GO:0002793, GO:0043043 | GO:1901652, GO:1901653 |
| sucrose | GO:0005986 | GO:0009744, GO:0106082 |
| trehalose | GO:0005992 | GO:0010353 |
| inositol | GO:0006021, GO:1900088, GO:1900089, GO:1900090 | GO:1902140, GO:1902141 |
| chitin | GO:0006031, GO:0032883 | GO:0010200, GO:0071323 |
| sorbitol | GO:0006061 | GO:0072708, GO:0072709 |
| ethanol | GO:0006115 | GO:0017036, GO:0045471, GO:0071361, GO:1901418 |
| camp | GO:0006171 | GO:0051591, GO:0071320 |
| cgmp | GO:0006182 | GO:0070305, GO:0071321 |
| l-arginine | GO:0006526 | GO:0010963, GO:0090467, GO:0097638, GO:1901042, GO:1902765, GO:1902826, GO:1902827, GO:1902828, GO:1903576, GO:1903577, GO:1905541, GO:1905542, GO:1905589 |
| glycine | GO:0006545, GO:0061536 | GO:0036233, GO:1900923, GO:1900924, GO:1900925, GO:1903804, GO:1905429, GO:1905430 |

| | | |
|---|---|---|
| proline | GO:0006561, GO:1902005, GO:1902006 | GO:0010238, GO:0071235, GO:1902825, GO:1902834, GO:1902835, GO:1902836, GO:1905647 |
| l-serine | GO:0006564 | GO:1903812 |
| octopamine | GO:0006589, GO:0061539 | GO:0071927, GO:2000128, GO:2000129, GO:2000130 |
| polyamine | GO:0006596, GO:0010967, GO:0170066 | GO:0140202 |
| fatty acid | GO:0000037, GO:0006633, GO:0042304, GO:0045717, GO:0045723 | GO:0070542, GO:0071398 |
| phosphatidylethanolamine | GO:0006646 | GO:1905711, GO:1905712 |
| cholesterol | GO:0006695, GO:0045540, GO:0045541, GO:0045542 | GO:0070723, GO:0071397 |
| ergosterol | GO:0006696, GO:0010895, GO:0032443, GO:0070452 | GO:1901625 |
| ecdysone | GO:0006697 | GO:0035075, GO:0071390 |
| bile acid | GO:0006699, GO:0032782, GO:0070857, GO:0070858, GO:0070859, GO:0120188, GO:0120189, GO:0120190 | GO:0038183, GO:1903412, GO:1903413 |
| progesterone | GO:0006701, GO:0042701, GO:2000182, GO:2000183, GO:2000184, GO:2000870, | GO:0032570 |

| | | |
|---|---|---|
| | GO:2000871, GO:2000872 | |
| androgen | GO:0006702, GO:0035935, GO:2000180, GO:2000834, GO:2000835, GO:2000836 | GO:2000825 |
| estrogen | GO:0006703, GO:0035937, GO:1904076, GO:1904077, GO:1904078, GO:2000861, GO:2000862, GO:2000863 | GO:0043627 |
| glucocorticoid | GO:0006704, GO:0031946, GO:0031947, GO:0031948, GO:0035933, GO:2000849, GO:2000850, GO:2000851 | GO:0051384 |
| mineralocorticoid | GO:0006705, GO:0035931, GO:2000855, GO:2000856, GO:2000857 | GO:0051385 |
| farnesol | GO:0006715 | GO:0097307, GO:0097308 |
| glutathione | GO:0006750, GO:1903786, GO:1903787, GO:1903788 | GO:0036347, GO:0072753, GO:0098709, GO:1901370 |
| atp | GO:0006754, GO:0006758, GO:0006759, GO:1904669, GO:2001169, GO:2001170, GO:2001171 | GO:0033198, GO:0071318 |
| heme | GO:0006783, GO:0070453, GO:0070454, | GO:1904334 |

| | | |
|---|---|---|
| | GO:0070455, GO:0097037 | |
| acetylcholine | GO:0008292, GO:0061526, GO:1905921, GO:1905922, GO:1905923 | GO:1905144, GO:1905145 |
| spermidine | GO:0008295, GO:1901304, GO:1901305, GO:1901307 | GO:0140203 |
| lipid | GO:0008610, GO:0046889, GO:0046890, GO:0051055, GO:0140353 | GO:0033993, GO:0071396 |
| pyridoxine | GO:0008615 | GO:1903075 |
| amino acid | GO:0008652, GO:0032973, GO:0044746, GO:0080143, GO:2000282, GO:2000283, GO:2000284 | GO:0010237, GO:0010958, GO:0043092, GO:0043200, GO:0044745, GO:0089718, GO:1902837 |
| methionine | GO:0009086 | GO:0044690, GO:0061431, GO:1903692, GO:1904640 |
| l-phenylalanine | GO:0009094, GO:0019274, GO:0019275 | GO:0140925 |
| l-leucine | GO:0009098, GO:2001276, GO:2001277, GO:2001278 | GO:0043201, GO:0060356, GO:0071233, GO:1903128, GO:1903129, GO:1903130, GO:1903801, GO:1905532, GO:1905533, GO:1905534 |
| l-valine | GO:0009099 | GO:0090468, GO:1903805 |
| glycoprotein | GO:0009101, GO:0010559, | GO:1904587, GO:1904588 |

| | | |
|---|---|---|
| | GO:0010560, GO:0010561 | |
| biotin | GO:0009102 | GO:0044756, GO:0070781, GO:0071296, GO:1901689, GO:1905135 |
| lipopolysaccharide | GO:0009103 | GO:0032496, GO:0071222 |
| vitamin | GO:0009110 | GO:0033273, GO:0071295 |
| nucleoside | GO:0009163 | GO:0180015 |
| pyrimidine ribonucleotide | GO:0009220 | GO:1905834, GO:1905835 |
| thiamine | GO:0009228, GO:0070623, GO:0070624, GO:0090180 | GO:0140125 |
| menaquinone | GO:0009234 | GO:0032572, GO:0071308 |
| cobalamin | GO:0009236 | GO:0033590, GO:0071297 |
| peptidoglycan | GO:0009252, GO:0009285 | GO:0032494, GO:0071224 |
| amine | GO:0009309 | GO:0014075 |
| putrescine | GO:0009446 | GO:1904585, GO:1904586 |
| gamma-aminobutyric acid | GO:0009449, GO:0014051, GO:0014052, GO:0014053, GO:0014054 | GO:0007214 |
| gibberellin | GO:0009686, GO:0010371, GO:0010372, GO:0010373 | GO:0009739 |
| abscisic acid | GO:0009688, GO:0010115, GO:0010116, GO:0090359 | GO:0009737 |
| cytokinin | GO:0009691 | GO:0009735 |
| ethylene | GO:0009693, GO:0010364, GO:0010365, GO:0010366, GO:0042456 | GO:0009723 |

| | | |
|---|---|---|
| jasmonic acid | GO:0009695, GO:0080141 | GO:0009753 |
| salicylic acid | GO:0009697, GO:0080142 | GO:0009751 |
| phenylpropanoid | GO:0009699 | GO:0080184, GO:1905546 |
| flavonoid | GO:0009813, GO:0009962, GO:0009963, GO:0009964 | GO:1905395, GO:1905396 |
| alkaloid | GO:0009821 | GO:0043279, GO:0071312 |
| auxin | GO:0009851, GO:0010600, GO:0010601 | GO:0009733 |
| amylopectin | GO:0010021 | GO:0044591, GO:1900523 |
| vitamin e | GO:0010189, GO:1904965, GO:1904966 | GO:0033197, GO:0071306 |
| brassinosteroid | GO:0010422, GO:0010423, GO:0016132, GO:2000488 | GO:0009741 |
| ketone | GO:0010566, GO:0042181 | GO:1901654, GO:1901655, GO:1990369 |
| vascular endothelial growth factor | GO:0010573, GO:0010574, GO:0010575, GO:1904046 | GO:0038084, GO:1900746, GO:1900747, GO:1900748 |
| norepinephrine | GO:0010700, GO:0010701, GO:0014061, GO:0042421, GO:0048243 | GO:0071873 |
| hydrogen peroxide | GO:0010728, GO:0010729, GO:0010730, GO:0050665 | GO:0042542, GO:0070301 |
| triglyceride | GO:0010866, GO:0010867, GO:0010868, GO:0019432 | GO:0034014, GO:0071401 |
| vitamin d | GO:0010957, GO:0042368, | GO:0033280, GO:0071305 |

| | | |
|---|---|---|
| | GO:0060556, GO:0060557 | |
| dopamine | GO:0014046, GO:0014059, GO:0033602, GO:0033603, GO:0042416, GO:1903179, GO:1903180, GO:1903181 | GO:1903350, GO:1903351 |
| epinephrine | GO:0014060, GO:0032811, GO:0032812, GO:0042418, GO:0048242 | GO:0071871 |
| pantothenate | GO:0015940, GO:0033317, GO:0033318 | GO:0044755, GO:0098717, GO:1901688 |
| carbohydrate | GO:0006093, GO:0016051, GO:0033231, GO:0043255 | GO:0009743, GO:0097319, GO:0098704 |
| triterpenoid | GO:0016104 | GO:1905836, GO:1905837 |
| sterol | GO:0016126, GO:0106118, GO:0106119, GO:0106120 | GO:0036314, GO:0036315 |
| glycoside | GO:0016138, GO:0016141 | GO:1903416 |
| antibiotic | GO:0017000 | GO:0046677, GO:0071236 |
| methylglyoxal | GO:0019242 | GO:0051595, GO:0097238 |
| rhamnose | GO:0019300 | GO:0032149 |
| mannose | GO:0019307 | GO:1905582, GO:1905583 |
| hexose | GO:0019319 | GO:0009746, GO:0140271 |
| dethiobiotin | GO:0019351 | GO:0044757, GO:1901690, GO:1905136 |
| acetate | GO:0019413 | GO:0010034, GO:0071311, GO:1901459 |
| mannitol | GO:0019593 | GO:0010555 |

| | | |
|---|---|---|
| l-ascorbic acid | GO:0019853, GO:2000082, GO:2000083 | GO:0033591, GO:0071298 |
| streptomycin | GO:0019872 | GO:0046679, GO:0071239 |
| peptide hormone | GO:0030072, GO:0090276, GO:0090277, GO:0090278 | GO:0043434 |
| insulin | GO:0030073, GO:0032024, GO:0046676, GO:0050796 | GO:0032868 |
| vasopressin | GO:0030103 | GO:1904116, GO:1904117 |
| bacteriocin | GO:0030152 | GO:0046678, GO:0071237 |
| growth hormone | GO:0030252, GO:0060123, GO:0060124, GO:0060125 | GO:0060416 |
| d-alanine | GO:0030632 | GO:0170048 |
| gonadotropin | GO:0032274, GO:0032276, GO:0032277, GO:0032278 | GO:0034698 |
| luteinizing hormone | GO:0032275, GO:0033684, GO:0033685, GO:0033686 | GO:0034699, GO:0042700 |
| hormone | GO:0032353, GO:0042446, GO:0046879, GO:0046883, GO:0046885, GO:0046886, GO:0046887, GO:0046888 | GO:0009725 |
| type i interferon | GO:0032479, GO:0032480, GO:0032481, GO:0032606, GO:0045351, GO:0072641 | GO:0034340, GO:0071357 |
| chemokine | GO:0032602, GO:0032642, | GO:1990868, GO:1990869 |

| | | |
|---|---|---|
| | GO:0032682, GO:0032722, GO:0042033, GO:0045073, GO:0045079, GO:0045080, GO:0050755, GO:0090195, GO:0090196, GO:0090197, GO:0090198 | |
| granulocyte macrophage colony-stimulating factor | GO:0032604, GO:0032645, GO:0032685, GO:0032725, GO:0042253, GO:0045423, GO:0045424, GO:0045425 | GO:0097012 |
| hepatocyte growth factor | GO:0032605, GO:0032646, GO:0032686, GO:0032726, GO:0048175, GO:0048176, GO:0048177, GO:0048178 | GO:0035728 |
| interferon-alpha | GO:0032607, GO:0032647, GO:0032687, GO:0032727, GO:0045349, GO:0045354, GO:0045355, GO:0045356, GO:0072642, GO:1902739, GO:1902740, GO:1902741 | GO:0035455, GO:0035457 |
| interferon-beta | GO:0032608, GO:0032648, GO:0032688, GO:0032728, GO:0035546, GO:0035547, GO:0035548, | GO:0035456, GO:0035458 |

| | | |
|---|---|---|
| | GO:0035549, GO:0045350, GO:0045357, GO:0045358, GO:0045359 | |
| type ii interferon | GO:0032609, GO:0032649, GO:0032689, GO:0032729, GO:0042095, GO:0045072, GO:0045077, GO:0045078, GO:0072643, GO:1902713, GO:1902714, GO:1902715 | GO:0034341, GO:0060332, GO:0071346 |
| interleukin-1 | GO:0032612, GO:0032652, GO:0032692, GO:0032732, GO:0042222, GO:0045360, GO:0045361, GO:0045362, GO:0050701, GO:0050704, GO:0050711, GO:0050716 | GO:0070555, GO:0071347 |
| interleukin-11 | GO:0032614, GO:0032654, GO:0032694, GO:0032734, GO:0042230, GO:0045363, GO:0045364, GO:0045365, GO:0072609, GO:0150169, GO:0150170, GO:0150171 | GO:0071105, GO:0071348 |
| interleukin-12 | GO:0032615, GO:0032655, GO:0032695, GO:0032735, GO:0042090, | GO:0070671, GO:0071349 |

| | | |
|---|---|---|
| | GO:0045075, GO:0045083, GO:0045084, GO:0072610, GO:2001182, GO:2001183, GO:2001184 | |
| interleukin-13 | GO:0032616, GO:0032656, GO:0032696, GO:0032736, GO:0042231, GO:0045366, GO:0045367, GO:0045368, GO:0072611, GO:2000665, GO:2000666, GO:2000667 | GO:0035962, GO:0035963 |
| interleukin-15 | GO:0032618, GO:0032658, GO:0032698, GO:0032738, GO:0042233, GO:0045372, GO:0045373, GO:0045374, GO:0072613 | GO:0070672, GO:0071350 |
| interleukin-17 | GO:0032620, GO:0032660, GO:0032700, GO:0032740, GO:0042235, GO:0045378, GO:0045379, GO:0045380, GO:0072615, GO:1905076, GO:1905077, GO:1905078 | GO:0097396, GO:0097398 |
| interleukin-18 | GO:0032621, GO:0032661, GO:0032701, GO:0032741, GO:0042241, GO:0045381, | GO:0070673, GO:0071351 |

| | | |
|---|---|---|
| | GO:0045382, GO:0045383, GO:0072616, GO:0150120, GO:0150121, GO:0150122 | |
| interleukin-2 | GO:0032623, GO:0032663, GO:0032703, GO:0032743, GO:0042094, GO:0045076, GO:0045085, GO:0045086, GO:0070970, GO:1900040, GO:1900041, GO:1900042 | GO:0070669, GO:0071352 |
| interleukin-21 | GO:0032625, GO:0032665, GO:0032705, GO:0032745, GO:0042238, GO:0045390, GO:0045391, GO:0045392, GO:0072619 | GO:0098756, GO:0098757 |
| interleukin-3 | GO:0032632, GO:0032672, GO:0032712, GO:0032752, GO:0042223, GO:0045399, GO:0045400, GO:0045401, GO:0072601 | GO:0036015, GO:0036016 |
| interleukin-4 | GO:0032633, GO:0032673, GO:0032713, GO:0032753, GO:0042097, GO:0042224, GO:0045402, GO:0045403, GO:0045404, GO:0072602, | GO:0070670, GO:0071353 |

| | | |
|---|---|---|
| | GO:0150133,<br>GO:0150134,<br>GO:0150135 | |
| interleukin-6 | GO:0032635,<br>GO:0032675,<br>GO:0032715,<br>GO:0032755,<br>GO:0042226,<br>GO:0045408,<br>GO:0045409,<br>GO:0045410,<br>GO:0072604,<br>GO:1900165,<br>GO:2000778 | GO:0070741,<br>GO:0071354 |
| interleukin-7 | GO:0032636,<br>GO:0032676,<br>GO:0032716,<br>GO:0032756,<br>GO:0042227,<br>GO:0045411,<br>GO:0045412,<br>GO:0045413,<br>GO:0072605,<br>GO:0150112,<br>GO:0150113,<br>GO:0150114 | GO:0098760,<br>GO:0098761 |
| interleukin-8 | GO:0032637,<br>GO:0032677,<br>GO:0032717,<br>GO:0032757,<br>GO:0042228,<br>GO:0045414,<br>GO:0045415,<br>GO:0045416,<br>GO:0072606,<br>GO:2000482,<br>GO:2000483,<br>GO:2000484 | GO:0098758,<br>GO:0098759 |
| interleukin-9 | GO:0032638,<br>GO:0032678,<br>GO:0032718,<br>GO:0032758,<br>GO:0042229,<br>GO:0045417,<br>GO:0045418, | GO:0071104,<br>GO:0071355 |

| | | |
|---|---|---|
| | GO:0045419,<br>GO:0072607 | |
| tumor necrosis factor | GO:0032640,<br>GO:0032680,<br>GO:0032720,<br>GO:0032760,<br>GO:0042533,<br>GO:0042534,<br>GO:0042535,<br>GO:0042536,<br>GO:1904467,<br>GO:1904468,<br>GO:1904469,<br>GO:1990774 | GO:0034612,<br>GO:0071356 |
| neurotrophin | GO:0032898,<br>GO:0032899,<br>GO:0032900,<br>GO:0032901 | GO:0038179 |
| nerve growth factor | GO:0032902,<br>GO:0032903,<br>GO:0032904 | GO:0038180,<br>GO:0051394,<br>GO:1990089 |
| isoquinoline alkaloid | GO:0033075 | GO:0014072,<br>GO:0071317 |
| raffinose | GO:0033529,<br>GO:1900091,<br>GO:1900092,<br>GO:1900093 | GO:0097403,<br>GO:1901545 |
| catecholamine | GO:0033604,<br>GO:0033605,<br>GO:0042423,<br>GO:0050432,<br>GO:0050433 | GO:0071869 |
| type iii interferon | GO:0034343,<br>GO:0034344,<br>GO:0034345,<br>GO:0034346,<br>GO:0072644 | GO:0034342,<br>GO:0071358 |
| cortisol | GO:0034651,<br>GO:0043400,<br>GO:0051462,<br>GO:0051463,<br>GO:0051464,<br>GO:2000064,<br>GO:2000065,<br>GO:2000066 | GO:0051414 |

| | | |
|---|---|---|
| vitamin a | GO:0035238 | GO:0033189, GO:0071299 |
| parathyroid hormone | GO:0035898, GO:2000828, GO:2000829, GO:2000830 | GO:0071107 |
| steroid hormone | GO:0035929, GO:0090030, GO:0090031, GO:0090032, GO:0120178, GO:2000831, GO:2000832, GO:2000833 | GO:0048545 |
| corticosterone | GO:0035934, GO:2000852, GO:2000853, GO:2000854 | GO:0051412 |
| testosterone | GO:0035936, GO:0061370, GO:2000224, GO:2000225, GO:2000843, GO:2000844, GO:2000845 | GO:0033574 |
| estradiol | GO:0035938, GO:2000864, GO:2000865, GO:2000866 | GO:0032355 |
| dehydroepiandrosterone | GO:0035942, GO:2000840, GO:2000841, GO:2000842 | GO:1903494, GO:1903495 |
| melanocyte-stimulating hormone | GO:0036160 | GO:1990680 |
| macrophage colony-stimulating factor | GO:0036301, GO:1901256, GO:1901257, GO:1901258 | GO:0036005, GO:0038145, GO:1902226, GO:1902227, GO:1902228, GO:1903971 |
| sodium ion | GO:0036376, GO:0071436, GO:0098667, GO:1903273, GO:1903274, GO:1903275, | GO:0097369, GO:0098719, GO:1903782, GO:1903783, GO:1903784, GO:1990118 |

| | | |
|---|---|---|
| | GO:1903276, GO:1903277, GO:1903278 | |
| bmp | GO:0038055, GO:1900144, GO:2001284, GO:2001285 | GO:0008101, GO:0030509, GO:0030510, GO:0030513, GO:0030514, GO:0071772, GO:0090097, GO:0090098, GO:0090099 |
| nicotine | GO:0042179 | GO:0035094, GO:0071316 |
| vitamin k | GO:0042371 | GO:0032571, GO:0071307 |
| phylloquinone | GO:0042372 | GO:0032573, GO:0071309 |
| long-chain fatty acid | GO:0042759 | GO:0010746, GO:0010747, GO:0010748, GO:0015911 |
| pheromone | GO:0042811 | GO:0019236, GO:0071444 |
| vitamin b6 | GO:0042819 | GO:0034516, GO:0071304 |
| pyridoxal | GO:0042821 | GO:0140204 |
| l-alanine | GO:0042852 | GO:1904273 |
| mycotoxin | GO:0043386 | GO:0010046, GO:0036146 |
| corticotropin-releasing hormone | GO:0043396, GO:0043397, GO:0051465, GO:0051466 | GO:0043435, GO:1900011 |
| alkane | GO:0043447, GO:1901577, GO:1901578, GO:1901579 | GO:1902778, GO:1902779, GO:1990373 |
| puromycin | GO:0043638 | GO:1905794, GO:1905795 |
| tetracycline | GO:0043644 | GO:0072746, GO:1901326 |
| macrophage migration inhibitory factor | GO:0044807 | GO:0035691, GO:2000446, GO:2000447, GO:2000448 |

| | | |
|---|---|---|
| adenine | GO:0046084, GO:0061934 | GO:0061488, GO:0098702 |
| guanine | GO:0046099 | GO:0061489, GO:0098710 |
| uracil | GO:0046107 | GO:0098721, GO:1902431, GO:1905529, GO:1905530, GO:1905531 |
| alcohol | GO:0046165, GO:1902930, GO:1902931, GO:1902932 | GO:0097305, GO:0097306, GO:1901421, GO:1990335 |
| methanol | GO:0046169 | GO:0033986, GO:0071405 |
| aldehyde | GO:0046184 | GO:0110096 |
| acetaldehyde | GO:0046186 | GO:1905640, GO:1905641 |
| cyanide | GO:0046202 | GO:1903927, GO:1903928 |
| toluene | GO:0046252 | GO:1901424, GO:1901456 |
| formaldehyde | GO:0046293 | GO:1904404, GO:1904405 |
| disaccharide | GO:0046351 | GO:0034285 |
| butyrate | GO:0043439, GO:0046358 | GO:1903544, GO:1903545 |
| monosaccharide | GO:0046364 | GO:0034284 |
| galactose | GO:0046369 | GO:0140425 |
| fructose | GO:0046370 | GO:0009750, GO:0032445, GO:1990539 |
| ceramide | GO:0046513, GO:1900060, GO:2000303, GO:2000304 | GO:0106096, GO:0106097 |
| tetrahydrofolate | GO:0046654 | GO:1904481, GO:1904482 |
| folic acid | GO:0046656 | GO:0051593, GO:0071231 |
| formic acid | GO:0046721 | GO:1901425, GO:1901462 |
| follicle-stimulating hormone | GO:0046880, GO:0046881, | GO:0032354, GO:0042699 |

| | | |
|---|---|---|
| | GO:0046882, GO:0046884 | |
| arachidonate | GO:0050482, GO:0090237, GO:0090238, GO:1900139 | GO:1904550, GO:1904551 |
| cocaine | GO:0050799 | GO:0042220, GO:0071314 |
| pullulan | GO:0051677 | GO:0044592, GO:1900520 |
| l-proline | GO:0055129 | GO:1903809, GO:1904271, GO:1905735, GO:1905736, GO:1905737 |
| copper ion | GO:0060003 | GO:0015678, GO:0046688, GO:0071280, GO:0098705, GO:1902861 |
| tyramine | GO:0061545, GO:1901695 | GO:0071928, GO:2000131, GO:2000132, GO:2000133 |
| l-glutamine | GO:0062132, GO:0062133, GO:0062134, GO:1901704 | GO:0036229, GO:1901034, GO:1901035, GO:1901036, GO:1903803, GO:1904844, GO:1904845 |
| glucagon | GO:0070091, GO:0070092, GO:0070093, GO:0070094 | GO:0033762 |
| somatostatin | GO:0070253, GO:0090273, GO:0090274, GO:0090275 | GO:0038170 |
| lipoteichoic acid | GO:0070395 | GO:0070391, GO:0071223 |
| prolactin | GO:0070459, GO:1902721, GO:1902722 | GO:0038161, GO:1902211, GO:1902212, GO:1902213, |

| | | |
|---|---|---|
| | | GO:1990637, GO:1990646 |
| thyroid-stimulating hormone | GO:0070460, GO:2000612, GO:2000613, GO:2000614 | GO:0038194 |
| hydrogen sulfide | GO:0070814, GO:1904826, GO:1904827, GO:1904828 | GO:1904880, GO:1904881 |
| l-asparagine | GO:0070981 | GO:0090469, GO:1903811 |
| l-methionine | GO:0071265 | GO:1903813, GO:1905544, GO:1905624, GO:1905625, GO:1905626 |
| homocysteine | GO:0071268 | GO:1905374, GO:1905375 |
| transforming growth factor beta | GO:0038044, GO:0071604, GO:0071634, GO:0071635, GO:0071636, GO:2001201, GO:2001202, GO:2001203 | GO:0071559 |
| chemokine (c-c motif) ligand 5 | GO:0071609, GO:0071649, GO:0071650, GO:0071651 | GO:0035689 |
| granulocyte colony-stimulating factor | GO:0071611, GO:0071655, GO:0071656, GO:0071657 | GO:0038158, GO:1990638, GO:1990643 |
| purine-containing compound | GO:0072522 | GO:0014074, GO:0071415 |
| interleukin-32 | GO:0072637, GO:0072638, GO:0150188, GO:0150189, GO:0150190, GO:0150191, GO:0150192, GO:0150193, GO:0150194 | GO:0097395, GO:0097397 |

| | | |
|---|---|---|
| fibroblast growth factor | GO:0090269, GO:0090270, GO:0090271, GO:0090272 | GO:0071774 |
| platelet-derived growth factor | GO:0090360, GO:0090361, GO:0090362 | GO:0036119 |
| l-glutamate | GO:0097054 | GO:0002036, GO:0002037, GO:0002038, GO:0098712, GO:1900920, GO:1900921, GO:1900922, GO:1902065, GO:1903802, GO:1905232, GO:1990123 |
| leukotriene b4 | GO:0097251 | GO:1905389, GO:1905390 |
| morphine | GO:0097295 | GO:0043278, GO:0071315 |
| chemokine (c-c motif) ligand 19 | GO:0097388 | GO:0038115 |
| chemokine (c-c motif) ligand 21 | GO:0097389 | GO:0038116 |
| chemokine (c-x-c motif) ligand 12 | GO:0097390 | GO:0038146 |
| potassium ion | GO:0071435, GO:0097623, GO:0098668, GO:1902302, GO:1902303, GO:1902304, GO:1903764, GO:1903765, GO:1903766 | GO:0010107, GO:0035864, GO:0035865, GO:1903287, GO:1903288, GO:1990573 |
| histidine | GO:0120213, GO:0120214, GO:0120215 | GO:0071232, GO:0080052 |
| manganese ion | GO:0140048 | GO:0010042, GO:0071287 |
| fluoride | GO:0140116 | GO:1902617, GO:1902618 |
| mycophenolic acid | GO:0140722 | GO:0071505, GO:0071506 |
| benzene | GO:1900997 | GO:1901423, GO:1901453 |

| | | |
|---|---|---|
| erythromycin | GO:1901115 | GO:0072743, GO:1901323 |
| ether | GO:1901503 | GO:0045472, GO:0071362 |
| strigolactone | GO:1901601 | GO:1902347, GO:1902348 |
| calcium ion | GO:1901660, GO:1905912, GO:1905913, GO:1905914, GO:1990034 | GO:0051592, GO:0071277, GO:0098703, GO:1905664, GO:1905665, GO:1905947, GO:1905949, GO:1990035 |
| cannabinoid | GO:1901696 | GO:0038171 |
| l-isoleucine | GO:1901705 | GO:0090476, GO:0090477, GO:1903806 |
| quercetin | GO:1901734 | GO:1905235, GO:1905236 |
| beta-carotene | GO:1901812 | GO:1905387, GO:1905388 |
| astaxanthin | GO:1901815 | GO:1905217, GO:1905218 |
| isobutanol | GO:1901961 | GO:1902665, GO:1990337 |
| l-dopa | GO:1903185, GO:1903195, GO:1903196, GO:1903197 | GO:1904473, GO:1904474 |
| glyoxal | GO:1903191 | GO:0036471 |
| reactive oxygen species | GO:1903409, GO:1903426, GO:1903427, GO:1903428 | GO:0000302, GO:0034614, GO:1901033 |
| iron ion | GO:1903414, GO:1903988 | GO:0010039, GO:0071281, GO:0097459, GO:0097460, GO:0098707, GO:0098711, GO:1903989, GO:1903990, GO:1903991, GO:1904438, |

| | | |
|---|---|---|
| | | GO:1904439, GO:1904440 |
| endothelin | GO:1904470, GO:1904471, GO:1904472, GO:1990775 | GO:1990839, GO:1990859 |

**Appendix D.** Supplementary figures showing representative interaction terms with approximately zero or negative causal enrichment scores (CES) across four datasets.

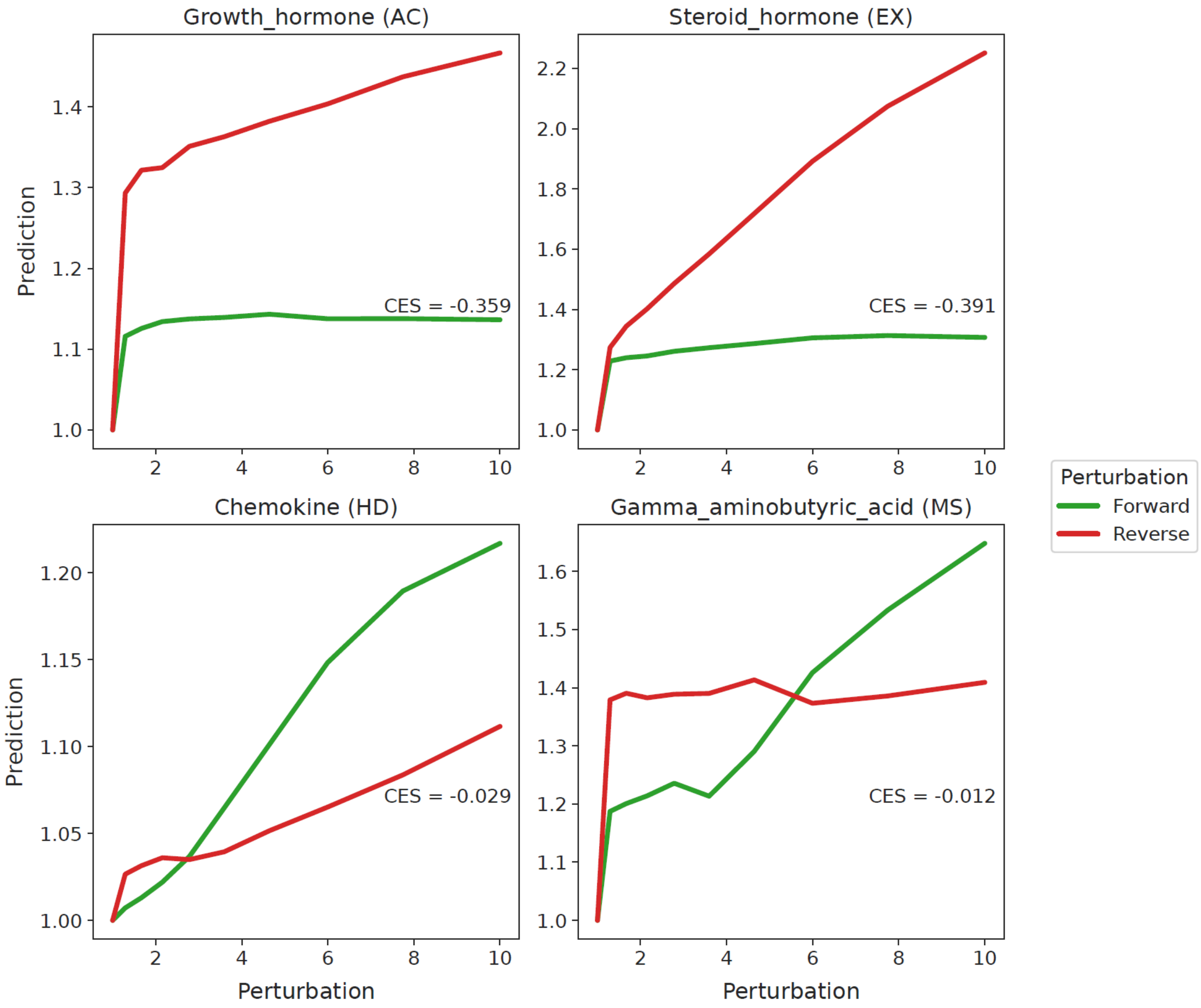
